# Optogenetic serotonin transmission in the prefrontal cortex is restrained by activity-upregulated voltage-gated potassium channels

**DOI:** 10.64898/2026.11.23.768547

**Authors:** Haushe Suganthan, Nikoo Ghafouri, Saige K. Power, Liz Rajesh, Claire L. Richter Gorey, Sridevi Venkatesan, Evelyn K. Lambe

**Author notes:** Corresponding author: Evelyn K. Lambe, Ph.D. Department of Physiology, Temerty Faculty of Medicine, University of Toronto 1 King’s College Circle Toronto ON, Canada M5S 1A8. Equally-contributing second authors (Nikoo Ghafouri, Saige K. Power). Equally-contributing senior authors (Sridevi Venkatesan, Evelyn K. Lambe).

## Abstract

Mood and anxiety are highly sensitive to the manipulation of serotonin receptors, but investigating relevant endogenous transmission with optogenetics has proved unexpectedly challenging. Here, we tested the reliability of optogenetic serotonin signalling in the prefrontal cortex, recording target cell electrophysiological responses to the stimulation of serotonin axons in brain slices from adult transgenic mice. The first train of optogenetic stimuli evoked the greatest response, which dropped to half on a subsequent train, minutes later. Focusing on the voltage-gated potassium channels classically involved in axon shunting, we found that serotonin neurons express a diversity of Shaker subunits, including the one associated with use-dependent functional upregulation. Blocking these channels with dendrotoxin-I rescued optogenetic response consistency. Reliable responses were also recapitulated with a low concentration of the Kv1 blocker, 4-aminopyridine. These interventions restore optogenetic outcomes to the naive level. Additional blockade of the high-voltage-sensitive Shaw channels expressed by serotonin neurons allowed further enhancement of optogenetic responses. Our findings suggest that Shaker and Shaw voltage-gated potassium channels work together to restrain prefrontal serotonin afferents in a use-dependent manner. Targeted blockers improve serotonergic transmission and may provide useful positive controls for the next steps in deciphering how our brains regulate mood and anxiety.

## Introduction

The prefrontal cortex plays an important role in higher-order tasks such as cognitive flexibility, emotional regulation, and executive function ^1,2^. Its excitability is regulated by serotonergic inputs from the dorsal and median raphe nuclei ^3–6^ in a manner thought to be essential for cognition, mood, and anxiety ^7–10^. Serotonin release in prefrontal cortex increases with stress ^11,12^, and disruptions in the serotonergic system can trigger neuropsychiatric conditions ^13–15^. Overall, serotonin neurotransmission in the prefrontal cortex is considered a major therapeutic target, with relevance for depression, anxiety, and panic disorders.

Most studies of serotonergic modulation have used exogenous ligands ^3,7,9,16^ to study this complex system with 14 receptor subtypes, including autoreceptors at the cell body level in the raphe nuclei and at axon terminals in the target regions ^17,18^. Yet the precise impact of *endogenous* serotonin signalling in the prefrontal cortex and other brain regions remains surprisingly enigmatic^19^.

With the advent of optogenetics, there has been a growing number of studies aimed at testing how the serotonin system affects cortical circuits and behaviour. This research has detected behavioural roles in cognitive flexibility and in promoting patience or persistence in probabilistic foraging ^20–27^. Significant behavioural effects have been observed with brief, infrequent optogenetic trains of 10-25 Hz to excite channelrhodopsin, and longer stimulations have proved effective at activating step-function opsins to facilitate release from endogenous activity ^23–25^. Brief ≥20 Hz excitation of red-shifted channelrhodopsin has been used to trigger endogenous release to be detected by new molecular sensors expressed in target regions ^29–33^, yet the serotonin increases detected have been small and overshadowed by profound subsequent decreases.

There are multiple effective strategies to label serotonin neurons and their axons via viral or transgenic approaches ^34–37,23^. These approaches allow robust optogenetic excitation at the level of the serotonin neurons in the dorsal raphe ^23,37–39^. However, serotonin appears challenging to release in its target regions *regardless* of whether the study is optogenetically stimulating in the raphe nucleus or in the terminal fields ^26,28,40,41^. The results of these *in vivo* and *ex vivo* studies are consistent with earlier work suggesting that serotonin release in target regions is tightly restricted^20,42–47^.

Limitations on serotonin release in the target regions would explain why pursuing optogenetic responses has demanded unorthodox approaches ^26,40,41,48^. Many studies have used high-copy viral approaches ^49^ together with *SERT-cre* knock-in mice, which have ∼50% less serotonin transporter and elevated serotonin levels ^35,50^. Yet even with these enhancements, the circuit, behavioural, and serotonin-sensing outcomes of optogenetically-released serotonin have been modest ^20–22,28,29,32,40^. These challenges contrast with the more straightforward optogenetic investigation of other targets, such as cholinergic or glutamatergic neurotransmission ^51–59^.

One potential explanation is that serotonin axons have distinctive properties that restrain their activation. Typically, excitatory opsins yield local depolarization but rely on the recruitment of threshold-activated sodium channels to relay the signal and trigger neurotransmitter release ^60,61^. However, in some types of axons, or under certain conditions, this recruitment could be prevented by local shunting conductances. Specific voltage-gated potassium (Kv) channels have been implicated in regulating terminal excitation threshold, firing properties, and levels of neurotransmitter release ^62–65^. Investigation in cell systems suggests that some Shaker Kv1 channels undergo use-dependent upregulation via an activation shift ^66^ that would further restrict subsequent transmitter release.

Here, we examine the reliability of optogenetic serotonergic stimulation to generate repeatable electrophysiological responses in prefrontal cortical slices. Floxed *Pet1-cre* mice have strong expression of channelrhodopsin in the serotonergic neurons of the dorsal and median raphe ^48^ and detectable optophysiological responses from axonal stimulation in the prefrontal cortex ^26,41^.

We demonstrate that the response elicited by a brief train of optogenetic stimuli at 20 Hz is strongest the first time it is delivered and declines substantially upon subsequent stimulation. Analysis of single-cell RNA sequencing (scRNA-seq) data indicates that serotonin neurons robustly express the Shaker Kv1.2 channel subunit, which is sensitive to use-dependent functional upregulation ^66^. Therefore, we tested the impact of blocking this channel on optogenetic replicability. Detecting a significant improvement, we extended this finding to a more accessible blocking strategy. Finally, we asked whether an additional class of high-voltage-sensitive shunting channels are recruited during optogenetic stimulation. Together, this work demonstrates that optogenetic stimulation of serotonergic transmission is restricted by two families of voltage-gated potassium channels.

## Results

### Prefrontal serotonergic optophysiological response diminishes upon repetition

To investigate the reliability of optogenetic stimulation of serotonin signalling, we optogenetically stimulate prefrontal cortical brain slices from adult *Pet1*cre^+^::Ai32^+/+^ mice (**Fig 1A**). These mice express channelrhodopsin in serotonergic neurons and their axons ^26,48^. We use whole cell recording from layer 5 pyramidal neurons to detect the serotonergic response, consistent with earlier characterization^26,27^. The first train of opto-stimulation (20 Hz, 5 s) elicits an outward current and this response is significantly reduced when the stimulus is repeated 5 minutes later (**Fig 1B, C**, a decline to 52 ± 6 % of the first response, paired t-test: *P*≤0.0001; n = 16 neurons from 8 mice). The layer 5 pyramidal neurons targeted for this investigation had the following intrinsic properties: membrane potential, −84 ± 1 mV; input resistance, 96 ± 12 MΩ; capacitance, 92 ± 6 pF; threshold, −52 ± 1 mV; and spike amplitude, 84 ± 2 mV (at rheobase).

**Figure 1.**
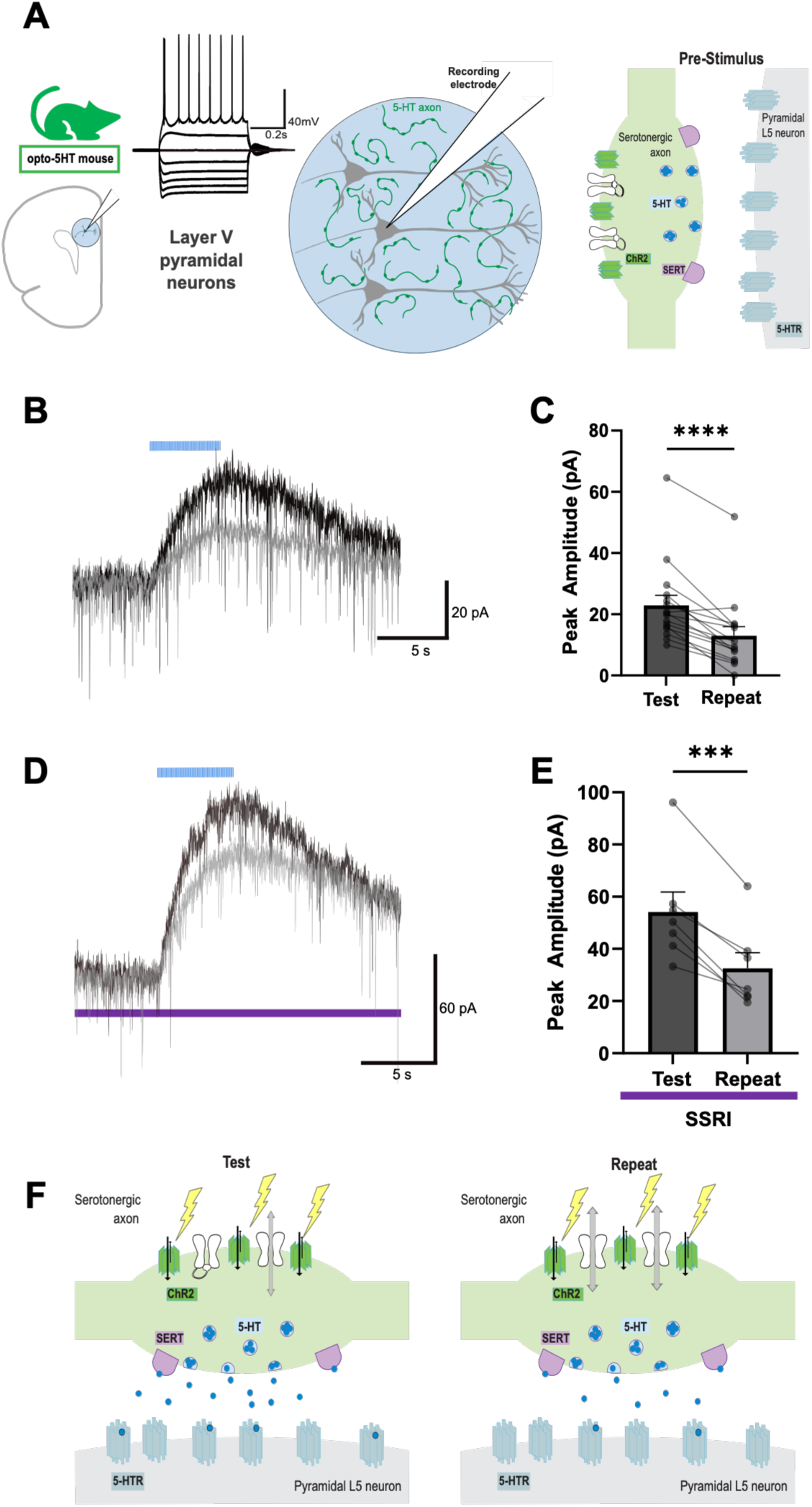
Repeated stimulation of serotonergic afferents yields a diminishing response. **A.** Schematic depicts the arrangement for optogenetic stimulation of serotonergic afferents in brain slices of prefrontal cortex from adult *Pet1cre^+^*::Ai32^+/+^ mice. Target responses are recorded in layer 5 pyramidal neurons, representative trace of a doublet-firing pyramidal neuron. Schematic of channels and receptors on the terminal and target sides before stimulation. **B**. Voltage-clamp recording of test (black) and repeat (gray) optogenetic stimulation of serotonergic afferents (blue bar, 20 Hz, 5 s), with the repeat delivered 5 min after the first-time stimulation. **C**. Scattergram shows the first-time and repeat responses are significantly different (t=6.806, df=15, p<0.0001, n=16, paired t-test). **D.** In presence of the selective serotonin reuptake inhibitor (SSRI), citalopram (1 µM, purple line), test and repeat optogenetic stimulation of serotonergic afferents. **E**. Scattergram shows the first-time and repeat responses in citalopram are significantly different (t=5.041, df=4, p=0.0073, n=5, paired t-test). **F.** Schematic representation of a serotonin axon terminal and its prefrontal neuron target during first-time (left) and repeat (right), illustrating the hypothesized change in optogenetic stimulation via a functional upregulation of ion channel shunting. Mean ± SEM, **** P < 0.0001.

Since serotonin reuptake is a potential source of activity-dependent suppression, we tested the first and repeat (test-repeat) responses in the presence of a selective serotonin reuptake inhibitor (SSRI) to block the serotonin transporter (SERT). As expected, acute application of citalopram (1 µM) substantially and significantly increased the amplitude of the first response compared to the previous experiment (control: 23 ± 3 pA, n = 16; citalopram: 54 ± 8, n = 9; unpaired t-test with Welch correction, t=3.754, df=8.299, P=0.0052). Yet, even with the much larger outward current on the first train, **Fig 1D, E** shows the significant suppression upon stimulus repetition (a decline to 59 ± 5% of first-time response; n = 7 neurons from 6 mice; paired t-test: *P*=0.0004).

### Serotonin neurons express Shaker voltage-gated potassium channel subunits

We hypothesized that the suppression on repeat stimulation occurs because of increased shunting conductance in serotonergic axon terminals. Accordingly, we turned to 10x single-cell RNA sequencing (scRNA-seq) data from the Allen Institute ^67^ of adult mice (n = 222 mice) to investigate expression of a class of terminal-trafficked, potassium channels in serotonergic neurons. We focused on the Shaker class since its Kv1.2-containing channels exhibit use-dependent functional upregulation ^66^. We visualized the expression patterns of the channel subunits, Kv1.1-Kv1.6 (encoded *Kcna1-Kcna6*) from the 10x scRNA-seq data ^67^. After filtering for serotonergic cells, we visualized genes of interest across a subsequent total of 1,469 cells. Since this study focuses on the serotonergic projections to the prefrontal cortex, Kvß2 (*Kcnab2*) was included for its relevance to Shaker subunit trafficking from the soma to the axon terminal ^68^. For context, we included the promoter used in the transgenic mice, Pet1 (*Fev*), and the serotonin transporter (*Slc6a4*), genes that contribute to the classification of serotonin identity ^67^.

This analysis demonstrates the presence of multiple Shaker family subunits in serotonin neurons, with stronger expression of *Kcna2* and *Kcna6* (**Fig 2A**), consistent with Kv1.2-containing channel upregulation in response to increasing neuronal activity ^66^. Allen data suggest 100% of these neurons express Kv1.2 channels together with potential partner Kv1.6, with Kv1.4 expressed in most and Kv1.5 in many. Consistent with terminal trafficking of Shaker family subunits, *Kcnab2* is robustly expressed. Also strongly expressed are Pet1 (*Fev*) and SERT (*Slc6a4*), markers of serotonin identity. This overall Shaker expression pattern is similar in female and male adult mice, with the sex segregated scRNA-seq data shown in **Suppl Fig S1**. Since Shaker channel subunits are expressed widely in the brain, we also show their expression in a few relevant cell classes alongside the serotonin cells (**Suppl Fig S2**).

**Figure 2.**
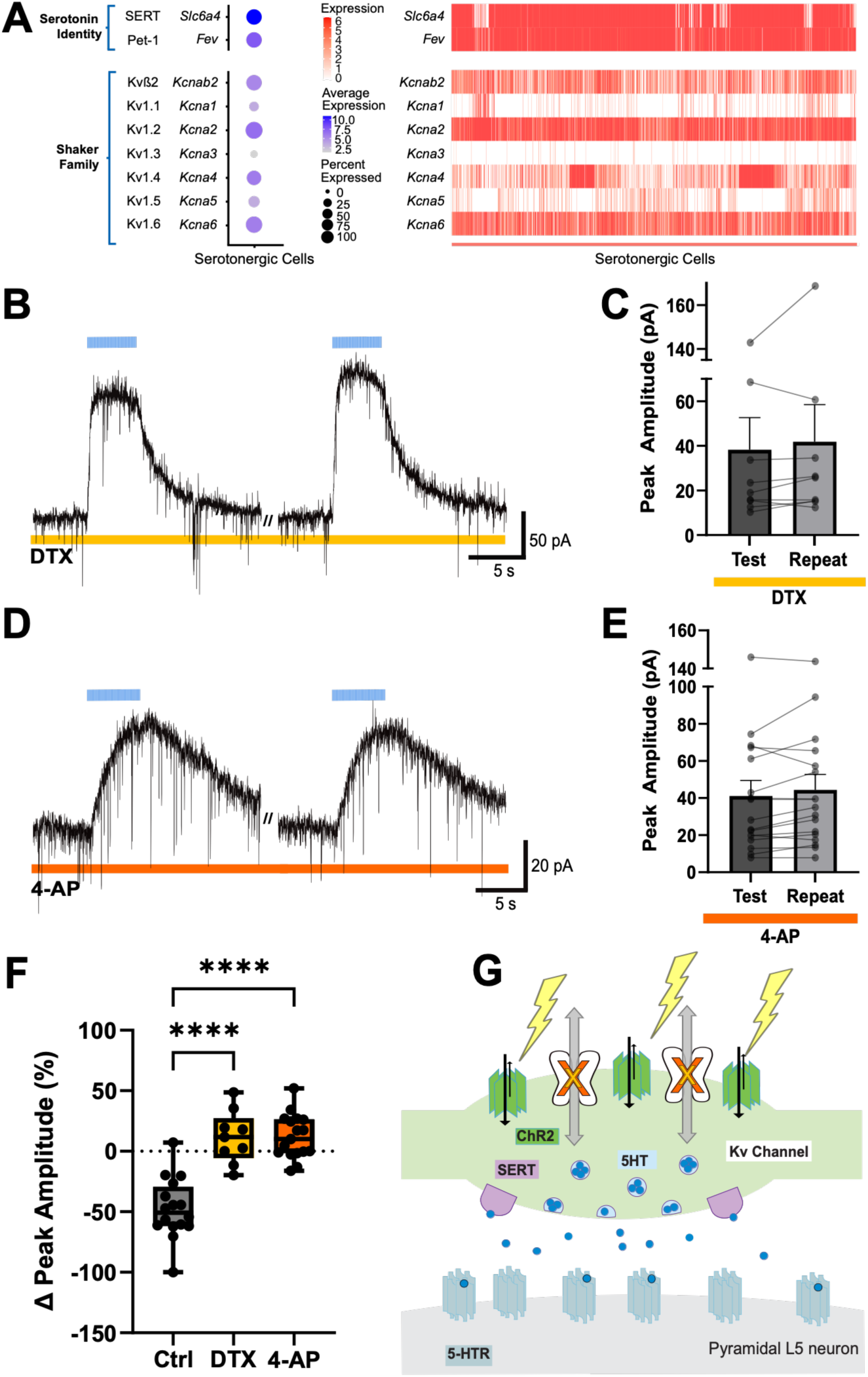
Serotonergic neurons express Shaker channel subunits and blocking these channels makes the optogenetic response repeatable. **A.** The dot plot (left) and the heat map (right) show that serotonergic neurons express the relevant subunits of the Shaker voltage-gated potassium channel family (Kv1), including the activity-upregulated Kv1.2, encoded by *Kcna2*, and Kvβ2 (*Kcnab2*), which are responsible for axonal trafficking of Shaker channels (Data from the Allen Brain Institute). **B.** Voltage-clamp recordings show the repeatability of the optogenetic response (blue bar, 20 Hz, 5 s) in the presence of dendrotoxin-I (DTX-I, 100 nM, yellow line), which blocks channels with a Kv1.1/1.2/1.6 subunit. **C.** Scattergram shows the repeatability of the test and repeat optogenetic responses in DTX-I (t=1.169, df=8, p=0.2762, n=9, paired t-test). Mean ± SEM **D.** Voltage-clamp recordings show the repeatability of the optogenetic response in the presence of a low concentration of 4-aminopyridine (4-AP, 5 μM, orange line). **E.** Scattergram shows the repeatability of the test and repeat optogenetic responses in 4-AP (t=1.915, df=16, p=0.0735, n=17, paired t-test). Mean ± SEM **F.** Graph contrasts the change in optogenetic response under different conditions: control, DTX-I, and 4-AP (one-way ANOVA: F _(2, 39)_ = 37.22, P<0.0001; Tukey’s multiple comparisons test: Control (Ctrl) vs. DTX-I: p<0.0001; Ctrl vs. 4-AP: p<0.0001; DTX-I vs. 4-AP: p=0.9984). **G.** Schematic of the serotonin axon terminal and its prefrontal neuron target during optogenetic stimulation during Shaker channel blockade. Box plot, **** P < 0.0001.

### Shaker channel blockade improves reliability of prefrontal optogenetic responses

To test whether blocking Shaker family Kv1 channels would rescue the reliability of the optogenetic response, we applied the Shaker-selective toxin dendrotoxin-I (DTX-I, 100 nM), which blocks channels containing at least one ^69^ of the subunits, Kv1.1/1.2/1.6. In its presence, opto-retest reliability is rescued (**Fig 2B,C**, n = 9 neurons from 5 mice; paired t-test: *P*= 0.2762). To extend the DTX-I finding, we tested the broader-spectrum blocker, 4-aminopyridine (4-AP; 50 µM) and found it also yields reliable optogenetic responses (test: 28.5 ± 5.9 pA; repeat: 26 ± 6 pA; paired t-test, t=1.899, df=3, P=0.1538; n=4 neurons from 3 mice); however, this higher concentration of 4-AP elicited substantial activity in the brain slice. Therefore, we tested lower concentrations of 4-AP that have been shown to exert significant effects in brain slice preparations ^70–72^. Strikingly, 4-AP (5 µM) fully rescued optogenetic test-repeat reliability (**Fig2D, E**, n = 17 neurons from 7 mice; paired t-test: *P*=0.0735, with a trend increase from 5-10 min 4-AP). Contrasting the change in peak amplitude (%) upon retest under control conditions and each type of Kv1 block, the differences are significant for both DTX-I and 4-AP (one-way ANOVA, Tukey’s multiple comparisons test: P<0.0001 and P<0.0001, respectively). Kv1 blockade resulted in minimal change in holding current (−6 ± 3 pA; one-sample t-test, t=1.803, df=20, P=0.0865; n=21 neurons from 10 mice). Test-retest suppression and rescue with Kv1 blockade are similar in male and female adult mice. **Suppl Fig S3** shows the sex-segregated data.

### Bidirectional confirmation that enhanced prefrontal optogenetic responses are serotonergic

To probe the serotonergic nature of the now-reliable optophysiological response, we measured its response to acute SSRI and to inhibition of serotonin receptors. In the presence of 4-AP, citalopram (1 µM) evoked an increase in peak amplitude not detected as significant (**Fig 3A-C**, 130 ± 12% n = 8 neurons from 4 mice; paired t-test: p=0.1908) together with a significant increase in the half- width of the optogenetic response (**Fig 3D**, 158 ± 14%; paired t-test: p=0.0045), consistent with mediation by release of endogenous serotonin. By contrast, application of a cocktail of serotonin receptor antagonists almost completely abolished the optophysiological response in the presence of 4-AP, peak amplitude was significantly reduced (**Fig 3E-G**, 25 ± 7% of the prior response, n = 11 neurons from 6 mice; paired t-test: p=0.0018) as well as half-width (**Fig 3H**, 77 ± 9%; paired t- test: p=0.0498). This bidirectional manipulation, with enhancement by SSRI and suppression by 5-HT receptor antagonists support the serotonergic nature of the optogenetic response in the presence of Shaker blockade.

**Figure 3.**
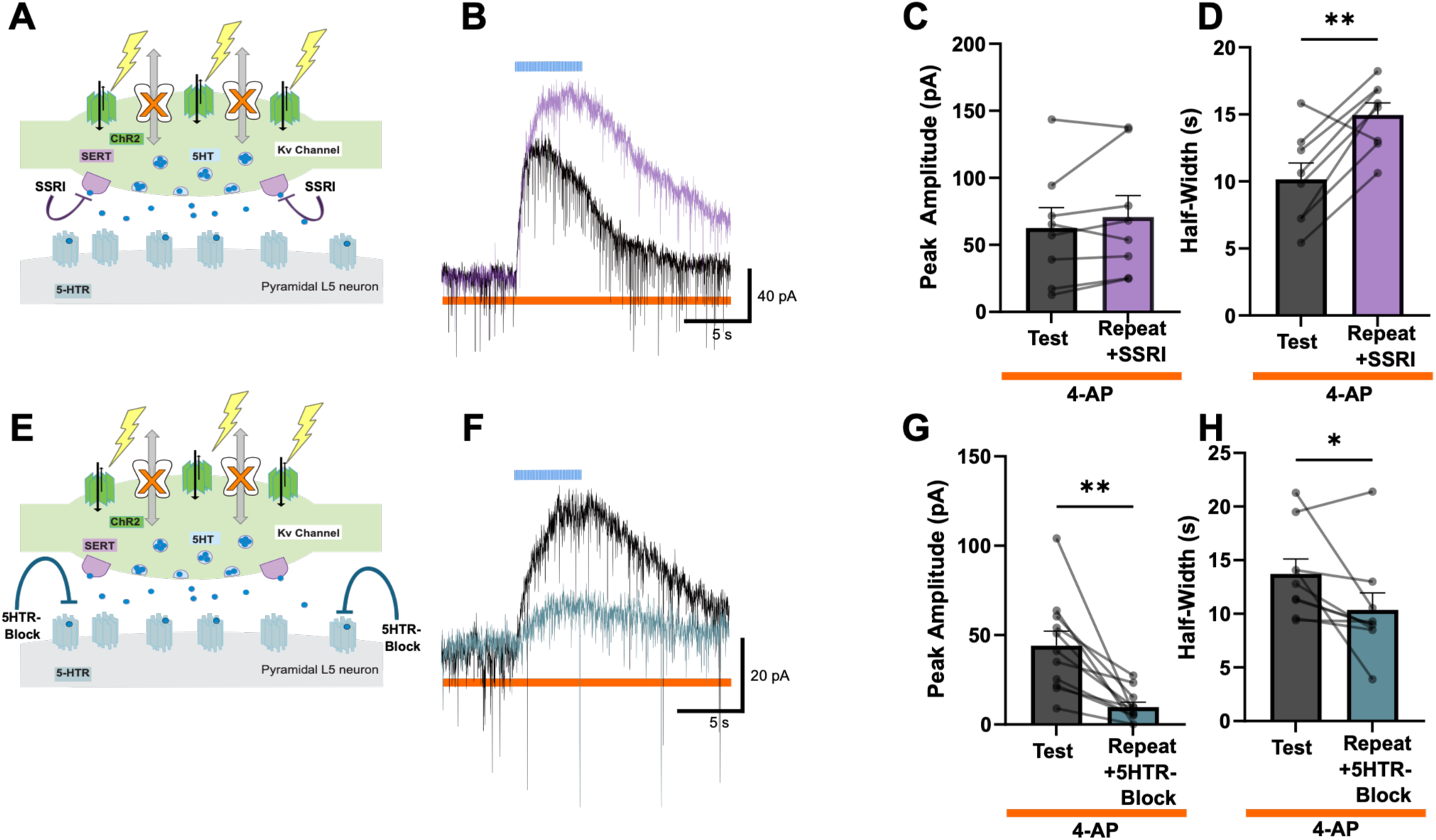
Reliable optophysiological responses during Shaker channel blockade are serotonergic. **A.** Schematic of serotonergic axon terminal and prefrontal neuron target, which illustrates Shaker channel block as well as inhibition of the 5-HT transporter with a selective serotonin reuptake inhibitor (SSRI). **B.** In the presence of 4-AP (5 μM, orange line), voltage-clamp recordings show the optogenetic response is enhanced by application of the SSRI, citalopram (1 μM, purple trace). Scattergrams show the SSRI effect on **C.** amplitude (t=1.448, df=7, p=0.1908, n=8, paired t-test) and **D.** half-width of the optogenetic response (t=4.119, df=7 p=0.0045, n=8, paired t-test). **E.** Schematic of the serotonergic terminal and prefrontal target illustrating Shaker channel block as well as an antagonist cocktail for 5-HT receptors (5-HTR Block, blue trace). **F.** In the presence of 4-AP, voltage-clamp recordings show the optogenetic response is suppressed by 5-HT receptor blockade. Scattergrams show the significant suppression by 5-HT receptor blockade of the **G.** amplitude (t=4.197, df=10, p=0.0018, n=11, paired t-test) and **H.** half-width of the optogenetic response (t=2.309, df=8 p=0.0498 n=9, paired t-test). Mean ± SEM, * *P* < 0.05, ** *P* < 0.01.

### Another voltage-gated potassium conductance restrains optogenetic serotonergic responses

To address the contribution of Shaker channels in suppressing the first-time (naive) optogenetic response, we evaluated how these responses compare to their subsequent rescue with Kv1 family blockade. When DTX-I or low dose 4-AP is applied after the initial test-repeat, **Fig 4A-E** shows that the optogenetic response is typically restored close to the level of the first-time response and only occasionally beyond (**Fig 4C, E**). Despite the likelihood of Shaker channel activation during the first-time optogenetic train, a significant increase beyond this response is not detected for either Shaker channel blocker (**Fig 4A-C**, DTX-I: n = 9 neurons from 5 mice; p=0.4942; **Fig 4D, E**, 4- AP: n = 14 neurons from 7 mice; p=0.6575, paired t-test). Therefore, we hypothesized that a high- voltage activated shunting conductance may be recruited in serotonergic terminals under these conditions.

**Figure 4.**
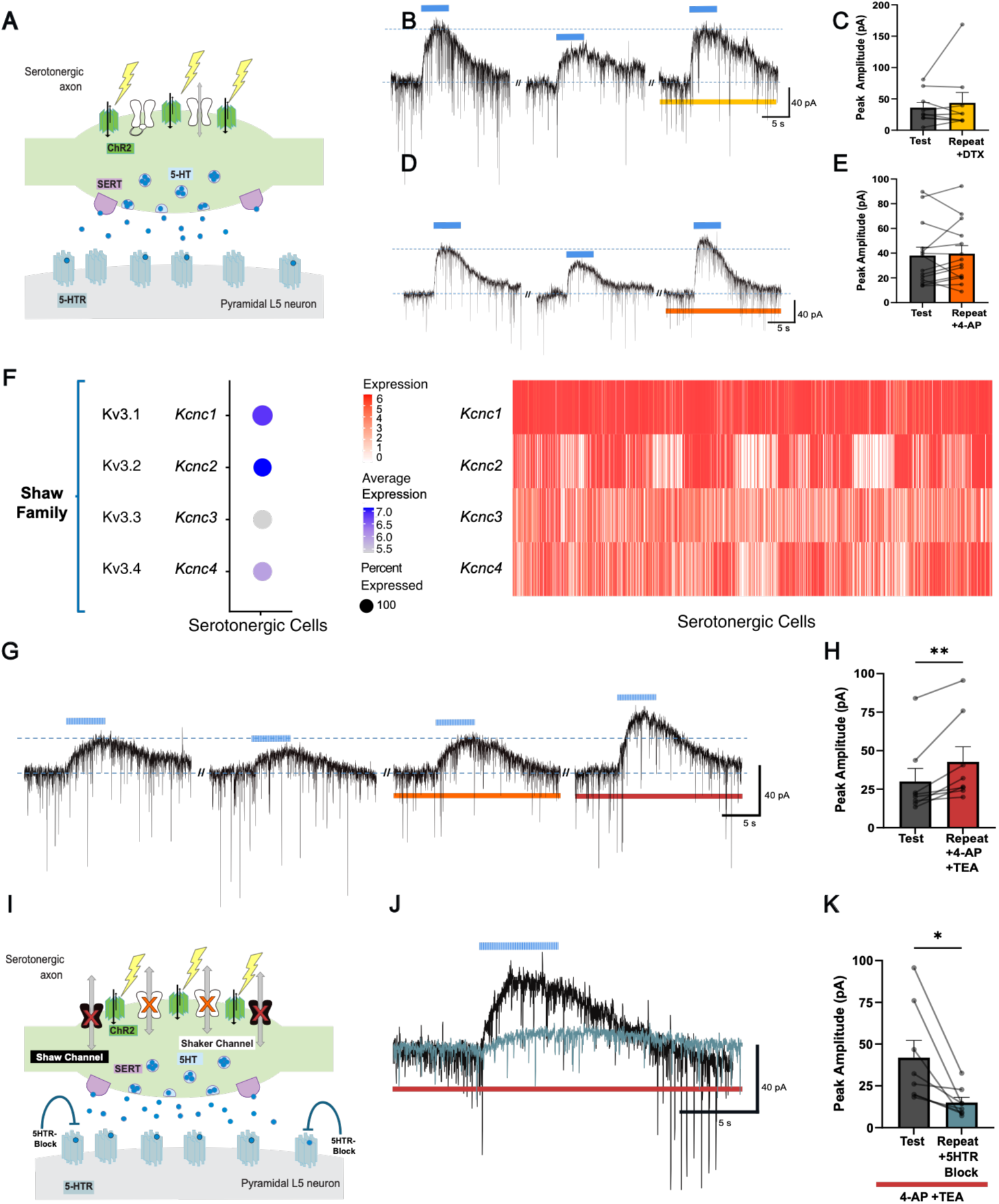
Shaker channel block restores naive optogenetic outcome but adding Shaw channel block yields greater serotonergic responses. **A.** Schematic of serotonergic axon terminal and prefrontal neuron target illustrating the first-time optogenetic stimulation. **B.** Voltage-clamp recordings of the response to optogenetic stimulation of serotonergic afferents (blue bar, 20 Hz, 5 s), with repeated stimulation every 5 min under control conditions and then in dendrotoxin-I (DTX- I, 100 nM, yellow line). **C.** Scattergram comparing the amplitudes of the first-time optogenetic response with those observed in DTX-I (t=0.3738, df=12, p=0.7151, n=13, paired t-test). **D.** Same experimental design with 4-aminopyridine (4-AP, 5 μM, orange line) to block Shaker channels. **E.** Scattergram comparing the amplitude of first-time optogenetic responses with those observed in 4-AP (t=0.4538, df=13, p=0.6575, n=14, paired t-test). **F.** The dot plot (left) and the heat map (right) show that serotonergic neurons express the relevant subunits of the Shaw high-voltage- activated potassium (Kv3) channel family, especially those encoded by *Kcnc1* (Data from the Allen Brain Institute). **G.** Voltage-clamp recordings with repeated optogenetic stimulation every 5 min under control conditions, with the addition of 4-AP, and with the further addition of tetraethylammonium (TEA, 1 mM, red line) to block Shaw family channels. **H.** Scattergram contrasting the amplitude of the first-time optogenetic responses with the significantly larger responses observed in 4-AP+TEA (t=3.663, df=7, p=0.0080, n=8, paired t-test). **I.** Schematic illustrating combined Shaker and Shaw channel blockade. **J.** Voltage-clamp recordings show a repeat of the enhanced optogenetic response in 4-AP+TEA (from example neuron in **G**) followed by its suppression by 5-HT receptor blockade (5HTR Block, blue trace). **K.** Scattergram demonstrating the significant suppression of the enhanced optogenetic responses in 4-AP+TEA by 5-HT receptor blockade (t=3.092, df=7, p=0.0175, n=8, paired t-test). Mean ± SEM, * *P* < 0.05, ** *P* < 0.01.

The Shaw family high-voltage-sensitive potassium channels have properties^73^ consistent with activation during the first-time optogenetic train as well as during the repeat trains in Shaker blockade. Analysis of serotonergic neurons in the scRNA-seq data shows robust expression of the Shaw channels, Kv3.1-3.4 (encoded by *Kcnc1-4)* (**Fig 4F**) and particularly of Kv3.1 (*Kcnc1*). The Shaw expression pattern is similar in females and males, and sex segregated data are shown in **Suppl Fig S4**. Since Shaw genes are widely expressed, we show their expression in relevant cell classes and serotonergic cells in **Suppl Fig S5** and tested the ability of Shaker and Shaw channel expression to differentiate serotonergic and cholinergic neuronal identity in **Suppl Fig S6**.

Since a diverse complement of Shaw channel channels are expressed by serotonin neurons, we investigated the impact of additionally blocking these channels on the optogenetic responses. In the presence of *combined* Shaw and Shaker blockade (tetraethylammonium, TEA, 1 mM; 4-AP, 5 µM), the optogenetic peak amplitudes reach 152 ± 14% of the first-time response, a significant increase (**Fig 4G-I**, p=0.0080, paired t-test; n = 8 neurons from 5 mice). By contrast, extended optogenetic responses remained similar to first-time responses when Kv1-only block was applied for longer (first-time 24.0 ± 5.7; extended Kv1 block 22.2 ± 5.0; paired t-test, t=0.2527, df =5, P=0.8016, n = 6). The enhanced optogenetic response with 4-AP+TEA remained consistent upon repetition (4-AP+TEA, 5 min: 39.0 ± 9.9 pA; 10 min: 41.9 ± 10.1 pA, paired t-test, t=1.647, df = 7, P=0.1435, n = 8). We did not detect a significant change in the holding current with the 4- AP+TEA blocker combination (−5 ± 4 pA; one-sample t-test, t=1.327, df=7, P=0.2261; n=8 neurons from 5 mice).

To probe the serotonergic nature of the enhanced optogenetic responses in Shaker and Shaw blockade, we tested their sensitivity to a cocktail of competitive serotonin receptor antagonists, observing strong and significant suppression (**Fig 4J-K**, 44 ± 8% of the prior response; n = 8 neurons from 5 mice; p=0.0175, paired t-test).

Taken together, these experiments reveal a pivotal and activity-sensitive role of voltage- gated potassium channels in dynamically suppressing serotonergic optogenetic responses.

## Discussion

Serotonergic afferents in the prefrontal cortex exert a more prominent response to first-time optogenetic stimulation. We hypothesized this diminishing phenomenon results from use- dependent functional upregulation of voltage-gated potassium channels in serotonergic afferents. Informed by scRNA-seq analysis, we targeted Kv1.1/1.2/1.6-containing channels. Dendrotoxin-I greatly improved test-repeat reliability of the optogenetic responses, an effect recapitulated by the nonspecific Shaker blocker, 4-AP, at a low concentration. The serotonergic nature of the reliable optogenetic responses was confirmed bidirectionally with serotonin reuptake inhibitor and serotonin receptor antagonists. Since the restoration appeared limited to the naive response amplitude, we hypothesized an additional role for the high-voltage-sensitive Shaw channels expressed by serotonin neurons in restraining optogenetic stimulation. Relieving shunting inhibition with the combination of Shaker and Shaw blockade significantly potentiated the optogenetic response beyond its original level, while retaining sensitivity to serotonin receptor block. This research identifies previously unsuspected mechanisms that limit optogenetic stimulation of endogenous serotonergic signalling, giving insight and tools to further investigate the native impact of serotonin on brain circuits and behaviour.

### New insight into voltage-dependent potassium channels in prefrontal serotonergic signalling

Our work suggests that Shaker channels play a dynamic role in the regulation of optogenetic serotonergic responses. The progressive nature of this suppression is a novel finding that builds on the identification of an activity-dependent upregulation of Kv1.2-containing Shaker channels through a hyperpolarizing shift in their activation curve ^66^. A parallel can be found in the inhibitory autoreceptor upregulation of axonal Kv1.2 channels to suppress electrically-evoked dopamine release ^74^. There is a long history of research on Shaker family control of neurotransmitter release ^64,75–78^, including optogenetic release of acetylcholine ^52,59,79^. Shaker blockade has predominantly been pursued in connection with optogenetic stimulation in the context of establishing whether such transmission is monosynaptic ^60,61,80–82^, where 4-AP was employed at more than an order of magnitude higher concentration (≥50µM) than what we discovered is sufficient to achieve reliable optogenetic stimulation of serotonin responses in the prefrontal cortex.

### Advances in this research and clinical implications

We have identified the progressive nature of the Shaker regulation of optogenetic serotonergic responses and elucidated the Shaw family channel recruitment. This work raises important questions about the potentially-distinct behavioural impact of “first-time” serotonin signalling after an interval. This scenario may have been detected in studies that found positive outcomes with optogenetic stimulations that were brief and given only infrequently ^20–22,26,28^. Our findings raise the question of how the Shaker and Shaw channel restraints on serotonin signalling are themselves regulated. Are these mechanisms always available for recruitment or are they selectively reduced or removed under certain behavioural or treatment conditions?

Intriguingly, there is a 4-AP analog, fampridine, which is an FDA-approved medicine for treatment of multiple sclerosis ^83–86^. In a small, randomized controlled trial, this 4-AP analog was found to exert positive effects on fatigue, cognition, and depression in patients who took it for two years ^87^. These positive effects were observed independent of changes in ambulatory status. This work suggests that blocking the voltage-gated potassium channels mostly highly sensitive to 4-AP improves human clinical symptoms associated with the serotonergic system ^15^.

It is increasingly appreciated that SSRIs are mechanistically complicated. They do not only inhibit the reuptake of serotonin but also exert blocking effects on Shaker and Shaw family channels (see review ^88^). In cell systems, SSRIs can block several different homomeric Shaker channels, as well as native Kv1 currents in hippocampal and cerebellar granule neurons ^89–95^. Shaw family Kv3.1 channels are also sensitive to blockade by SSRIs and related compounds ^89,96–98^. While this prior research has largely been pursued in cell systems, our findings raise the possibility of sensitive interactions between SSRIs and the native voltage-gated potassium channels which regulate serotonergic neurotransmission.

### Mechanistic caveats

To confirm involvement of the Shaker family, we selected the DTX-I toxin, which blocks Kv1.1/1.2/1.6 subunits. Of note, Shaker family use-dependent upregulation has been shown in cell systems to be conferred on its tetrameric channels containing even a single Kv1.2 subunit ^66^. We hypothesize the relevant Shaker channels may be complex heteromers based on subunit expression (**Fig 2**). Heteromultimeric channels containing Kv1.2, Kv1.4, and Kv1.6 would be consistent with the expression data, and cell systems research shows this configuration yields slowly-inactivating currents ^99,100^. However, limitations in the selectivity of the available toxins and blockers make it challenging to confirm the involvement of this specific heteromultimer.

The Shaw family, moreover, has strong expression in serotonin neurons, but there are even fewer selective choices for its blockade. We applied tetraethylammonium (**Fig 4**) at a concentration suggested to be selective ^73^ and at which its impact is lost in a Kv3.1/3.2 double-knockout mouse ^101^. Blood depressing substance-1 (BDS-1) has also been suggested to be selective towards Kv3.1/3.2/3.4 ^73^. However, BDS-1 also acts on sodium channels in a variety of neurons^102^, and slowing inactivation of voltage-gated sodium channels would complicate examination of the impact of Shaw channels on optogenetic serotonergic neurotransmission. Further work with Kv3 positive allosteric modulators may clarify the specific Shaw channels involved ^103,104^. although these experiments would likely be challenging to interpret.

Our work interrogates the impact of endogenous serotonin, optogenetically released, on the local circuitry of the mouse prefrontal cortex. This *ex vivo* focus does not permit us to examine the degree to which Kv1 and Kv3 channels in serotonin axons and terminals actively shunt action potentials which are initiated at the level of the raphe, whether by specific frequencies of optogenetic stimulation or by the more-variable behavioural firing ^8,39,105^. Serotonin appears challenging to release in its target regions *regardless* of whether the study is optogenetically stimulating in the raphe nucleus or in the terminal fields ^20–22,28,29,32,40,41^, this appears to be a promising avenue for future research.

### Significance and impact of the findings

Overall, our work demonstrates that optogenetic excitation of serotonin afferents in the prefrontal cortex is under complex, activity-dependent constraint by Shaker and Shaw voltage-gated potassium channels. These results identify important tools and insights towards further deciphering the regulation of the serotonergic system.

## Materials and Methods

### Animals

The University of Toronto Faculty of Medicine Animal Care Committee approved all experiments in accordance with the guidelines of the Canadian Council on Animal Care (protocol #20011621). Adult mice were used for all studies (age: P147 ± 17 postnatal days, mean ± SEM). Both male and female mice were included, and we show sex-disaggregated data for the major findings. We performed experiments on brain slices from transgenic crosses of *Pet1Cre*^+^ ^106^ and floxed Ai32 mice, as previously described and validated as serotonin-selective^26,27,48^. The mice were a kind gift from Dr. Mark Ansorge of Columbia University.

### Electrophysiological Recordings

Electrophysiology experiments were conducted in brain slices from adult male and female *Pet1*cre^+^::Ai32^+/+^mice. Coronal slices (400 µm thick) of prefrontal cortex were prepared using a Dosaka linear slicer (SciMedia) in 4°C sucrose artificial cerebrospinal fluid (aCSF) (254 mM sucrose, 10 mM D-glucose, 24 mM NaHCO_3_, 2 mM CaCl_2_, 2 mM MgSO_4_, 3 mM KCl, 1.25 mM NaH_2_PO_4_; pH 7.4). Slices were then put to recover for at least 2 hours in oxygenated (95% O2, 5% CO_2_) aCSF (128 mM NaCl, 10 mM D-glucose, 26 mM NaHCO_3_, 2 mM CaCl_2_, 2 mM MgSO_4_, 3 mM KCl, and 1.25 mM NaH_2_PO_4_; pH: 7.4) at 30°C before being used for whole-cell electrophysiological experiments. To continue to maintain synthesis of serotonin brain slices were recovered and recorded in the presence of 30 µM L-tryptophan ^107^.

For whole-cell patch clamp electrophysiology, brain slices were transferred to a BX51WI microscope (Olympus microscope, RRID:SCR_023069) and positioned in a chamber constantly perfused with oxygenated ACSF at 32°C. Layer 5 pyramidal neurons were patched based on their laminar localization and morphology in the prelimbic and cingulate regions of the medial prefrontal cortex. The recording electrodes (2-3 MΩ) were filled with intracellular recording solution composed of 120 potassium gluconate, 5 mM KCl, 2 mM MgCl_2_, 4 mM K_2_-ATP, 0.4 mM Na_2_-GTP, 10 mM disodium-phosphocreatine and 10 mM HEPES buffer and with pH adjusted to 7.3 using KOH. Recordings were obtained using an Axopatch200B amplifier (RRID:SCR_018866, Molecular Devices) at 20 kHz and low pass filtered at 3 kHz using pClamp (RRID:SCR_011323, Molecular Devices) and corrected for the liquid junction potential (14 mV). The neuronal firing pattern and characteristics were tested with steps in current-clamp before proceeding to optogenetic experiments in voltage-clamp at a holding potential of −75 mV.

### Optogenetics

Optogenetic stimulation was performed on layer 5 pyramidal neurons which are doublet-firing and have an H-current sag, as illustrated in the example in Fig 1A. This group has been suggested to be enriched for extratelencephalic neurons ^108–112^, but may include some intratelencephalic neurons ^113^. Extratelencephalic neurons have been implicated in stress response ^114^ and suggested to exhibit a relatively homogenous response to locally-applied serotonin ^115^. To excite channelrhodopsin- containing axonal fibers, optogenetic stimulation was delivered by blue light (470 nm) using a microscope mounted collimated LED (Thorlabs, Newton, New Jersey, United States) as follows: 10 ms light pulses in a 20 Hz train for 5 s. Optogenetic stimulus trains were separated by 5-min intervals, except for a subset of cells in **Fig 1D** with 10-min intervals.

### Pharmacology

Dendrotoxin-I (DTX-I, 100 nm, Alomone, 100 nM) was given to block Kv1.1/1.2/1.6-containing channels, and 4-aminopyridine (4-AP, Sigma, 5 µM, except one specified experiment at 50 µM) was used for blockade of Kv1.x channels. Tetraethylammonium (TEA, Sigma 1 mM) was given to block Kv3.x channels. The antagonist cocktail for serotonin receptors (5HTR block) included a combination of WAY100635 (100 nM, Tocris) to block 5-HT_1A_ receptors and SB699551 (20 µM, Tocris) to block 5-HT_5A_ receptors ^116^ and/or methiothepin (10 µM, Tocris), for a broader 5-HT receptor block. Citalopram (1 µM, Focus Biomolecules) was used to inhibit the serotonin transporter (SERT) in a subset of experiments.

### Inclusion and Exclusion Criteria

Our inclusion criteria for optogenetic experiments meant that we investigated layer 5 pyramidal neurons that were doublet-firing and had an H-current sag, as ascertained with current injections in current-clamp. Only one optophysiological series was performed per brain slice. The naive responses are the first time the brain slice received optogenetic stimulation. All datapoints from completed experiments were included in the analysis. Our exclusion criteria were loss of access or deterioration in cell health during the experiment (n=4). We included both male and female mice and show sex-disaggregated data for the major findings (**Suppl. Fig S1, S3**, and **S4**).

### Statistical Analysis

GraphPad Prism 11.0.2 (RRID:SCR_002798) was used for the statistical analyses and graphs for the optophysiology experiments. Statistical analyses included Student’s two-tailed paired and unpaired t-tests for two group comparisons. Two-tailed one sample tests were used for one group comparisons. One-way ANOVA with Tukey’s multiple comparison tests were used to contrast three groups. Power analyses for major comparisons were calculated (β=0.8, α=0.05) based on effect size in pilot experiments. Data were reported as mean ±SEM, except for the box plot with median and quartiles in **Fig 2F**. Significance was set at P<0.05.

### Transcriptomic Analysis

For our transcriptomic analysis of *Shaker* and *Shaw* potassium channel gene expression, we used the publicly available 10x scRNA-seq dataset ^67^ from the Allen Institute’s whole mouse-brain cell type atlas. We used the Seurat package (Seurat version 5.5.0, RRID:SCR_007322) ^117^ in R software (R version 4.6.0, RRID:SCR_001905) to subset the data for cells assigned with the Allen Institute’s ‘Sero’ label for serotonergic identity. Heat maps and dot plots of the resulting 1,469 cells were generated with Seurat DoHeatmap and DotPlot. Sex-segregated heat maps were generated after a further filtering serotonergic cells for male or female donor cells. Violin and ridge plots of the cholinergic and layer 5 glutamatergic neurons, in addition to the serotonergic neurons, were generated with Seurat’s VlnPlot and RidgePlot functions. The cholinergic cells were subset with the ‘Chol’ label (7,578 cells); cerebral cortex layer 5 intratelencephalic (L5 IT; 15,020 cells) and extratelencephalic (L5 ET; 5,710 cells) glutamatergic cells were subset by their respective transcriptomic subclasses (‘005 L5 IT CTX Glut’, ‘022 L5 ET CTX Glut’). Selection of these cell populations for contrast was motivated by their less complicated recruitment when targeted optogenetically ^51–59^. The cortical glutamatergic populations likely contain the layer 5 pyramidal neurons in the electrophysiological recordings. Differential gene expression analysis of voltage- gated potassium channel gene expression between serotonergic and cholinergic cells was performed using Seurat’s FindMarkers function with default parameters, except for test.use set to ‘roc’. The result was visualized as a dot plot using ggplot2 (RRID:SCR_014601).

## Acknowledgements & Funding

We thank Ms. Yao-Fang Tan for expert technical assistance. The research was funded by an NSERC Discovery Grant (EKL). HS was partially supported by a Postdoctoral Fellowship from the Department of Physiology. SKP received Canada Graduate Scholarship Doctoral and Postdoctoral Awards. CRG is supported by a Canada Graduate Scholarship Doctoral Award. NG and LR received University of Toronto Undergraduate Research Awards.

## Contributions

Conceptualization (HS, NG, SV, EKL); Investigation (HS, NG, SKP, LR, CRG); Analysis (HS, NG, SV); Visualization (HS, NG, SV); Writing - original draft (HS, NG); Writing - review and editing (all authors); Supervision (SV, EKL); Resources (EKL).

## Data Availability

All optophysiology datapoints supporting the findings of this study are displayed in scattergram format within the article and the supplementary files. The scRNA-seq data are publicly available from the Allen Brain Institute.

---

**Supplementary Figure S1.**
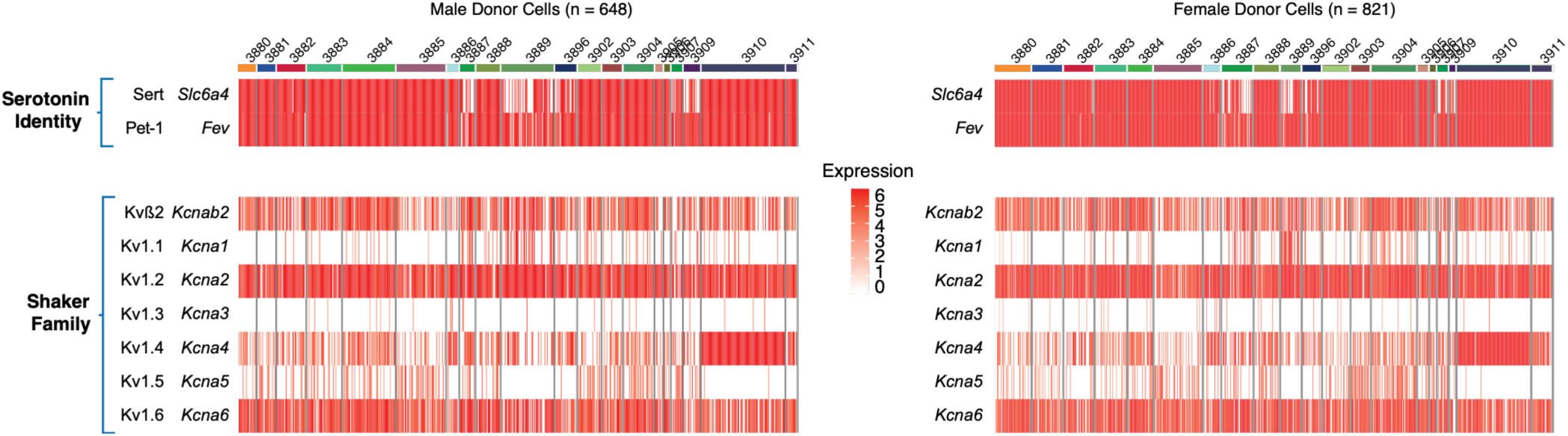
In both male and female mice, serotonergic neurons express Shaker voltage-gated potassium channel subunits. Heat maps of Shaker subunit expression across n = 648 male donor cells (left) and n = 821 female donor cells (right) sequenced from serotonergic dorsal raphe neurons. Cells are grouped by cluster identity. Heat maps show consistent cell cluster identities and patterns of Shaker subunit gene expression when disaggregated by sex, including the activity-upregulated Kv1.2 channel (*Kcna2*) and the Kvß2 (*Kcnab2*) subunit, which is responsible for axonal trafficking of Shaker channels. Data from the Allen Brain Institute.

**Supplementary Figure S2.**
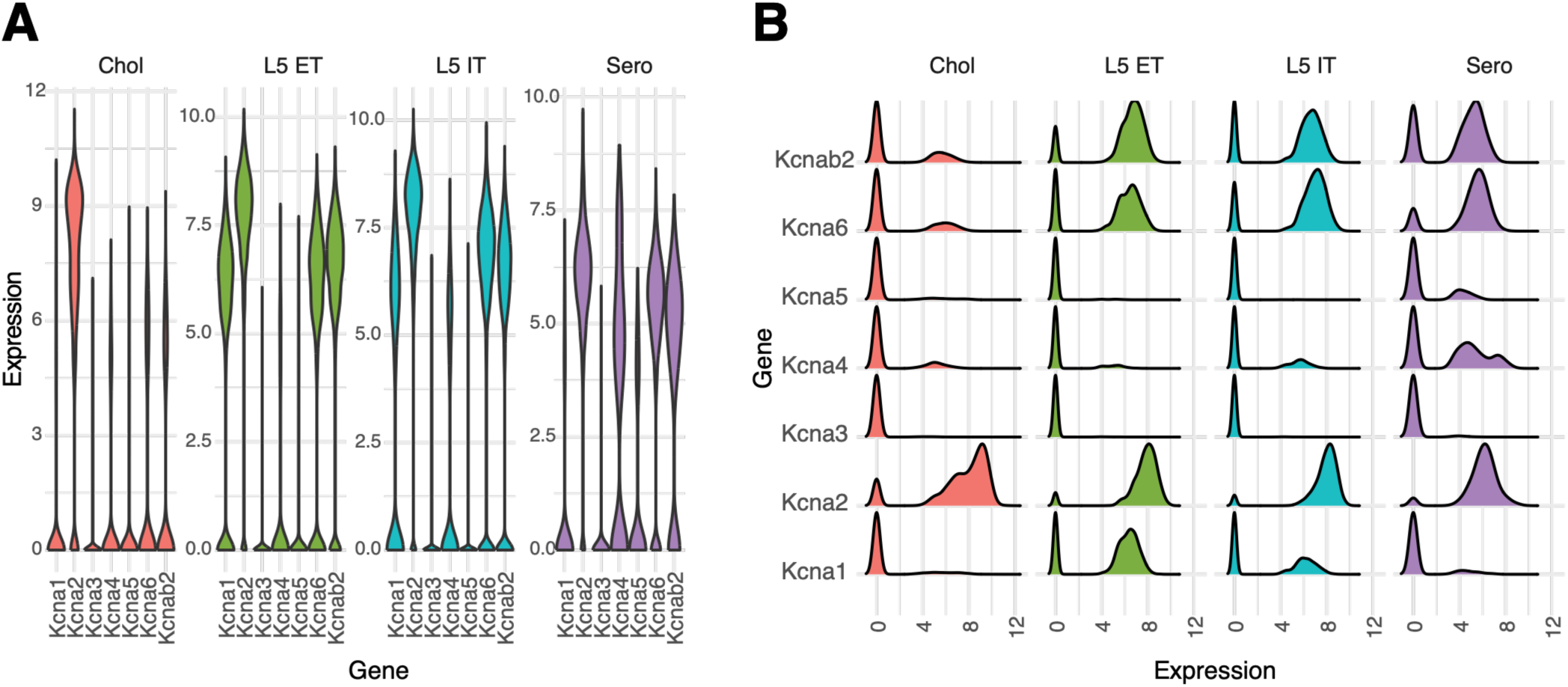
Expression of Shaker voltage-gated potassium channel subunits in some relevant cell classes. **A.** Violin plots and **B.** ridge plots of Shaker voltage-gated potassium channel subunit (Kv1) expression, including Kvß2 (*Kcnab2*), a subunit responsible for Shaker channel axonal trafficking. Expression is visualized across cholinergic cells (red), two subtypes of cortical glutamatergic layer 5 cells (L5 ET, green; L5 IT, blue), and serotonergic cells (purple). Data from the Allen Brain Institute.

**Supplementary Figure S3.**
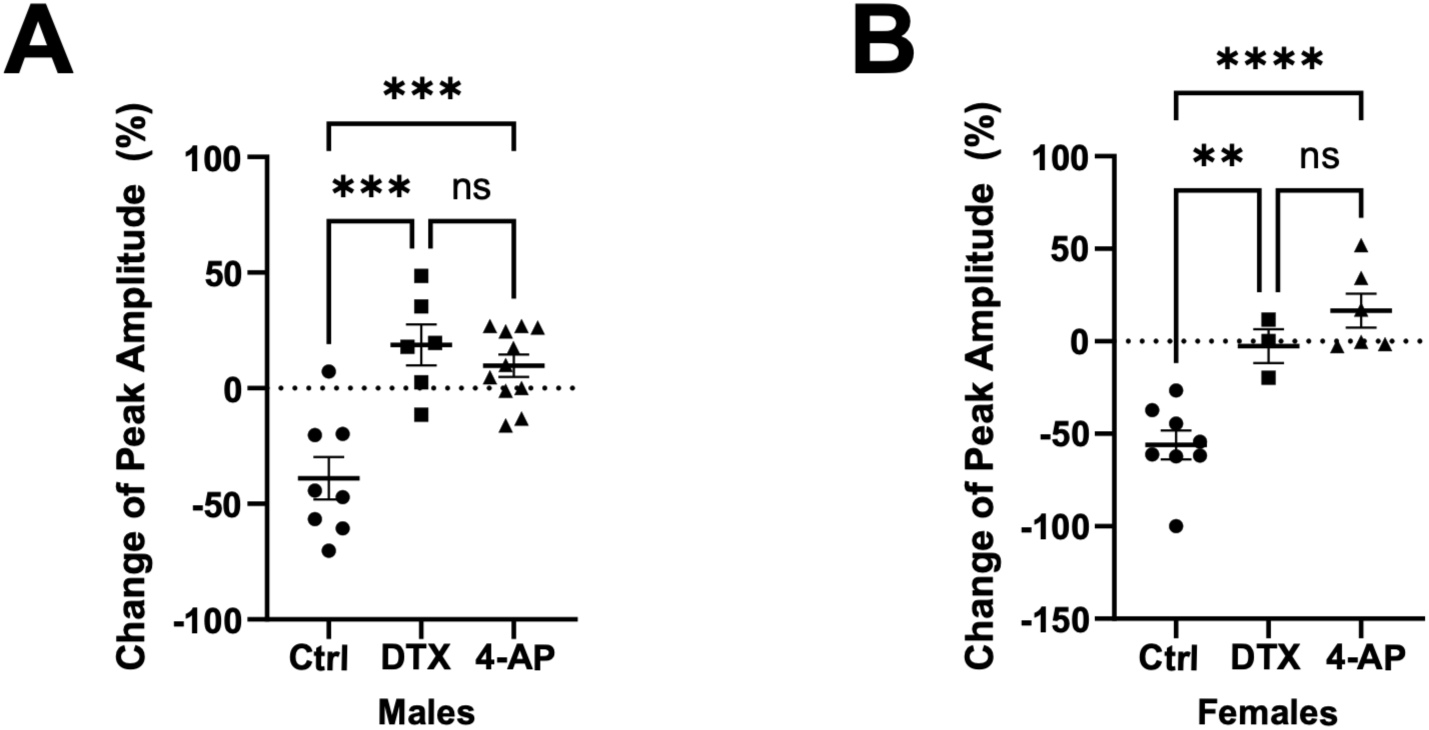
In both male and female mice, serotonergic responses diminish with repeat optogenetic stimulation under control conditions but are replicable under Kv1 blockade. Each graph contrasts the percent change in optogenetic response for test-repeat under three conditions: control (Ctrl), dendrotoxin-I (DTX-I), and 4-aminopyridine (4-AP). **A**. In males, there is a significant effect of condition (one-way ANOVA: F _(2, 22)_ = 17.09, P<0.0001; Tukey’s multiple comparisons test: Ctrl vs. DTX-I: p=0.0001; Ctrl vs. 4-AP: p=0.0001; DTX-I vs. 4-AP: p=0.6734. **B.** In females, there is a significant effect of condition (one-way ANOVA: F _(2, 14)_ = 20.93, P<0.0001; Tukey’s multiple comparisons test: Ctrl vs. DTX-I: p=0.0066; Ctrl vs. 4-AP: p<0.0001; DTX-I vs. 4-AP: p=0.4370). Mean ± SEM, ns *P* > 0.05, \*\**P* < 0.01,*** P < 0.001, **** P < 0.0001.

**Supplementary Figure S4.**
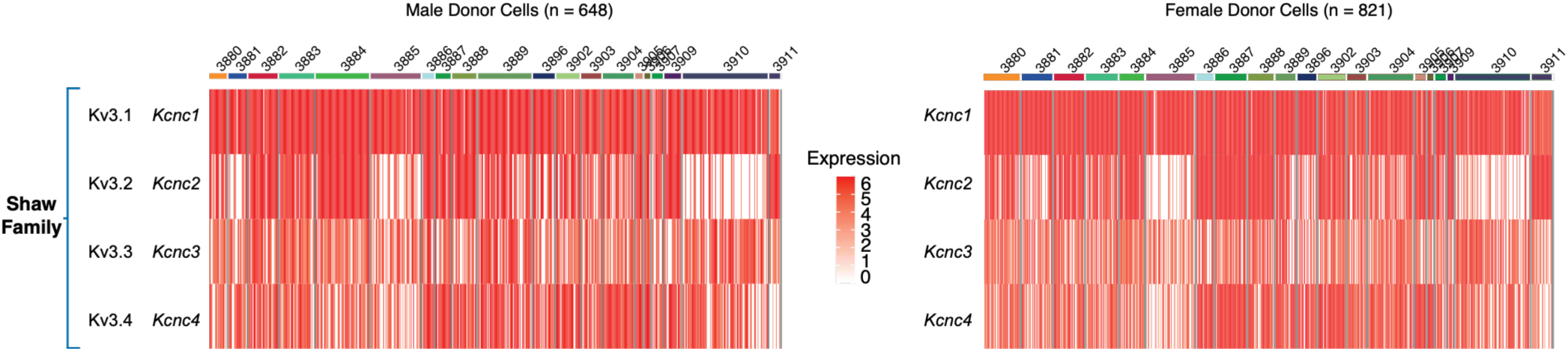
In both male and female mice, serotonergic neurons express Shaw voltage-gated potassium channel subunits. Heat maps of Shaw high-voltage-gated potassium channel family (Kv3) subunit expression across n = 648 male donor cells (left) and n = 821 female donor cells (right) sequenced from serotonergic raphe neurons. Cells are grouped by cluster identity. Heat maps show consistent cell cluster identities and patterns of Shaw subunit gene expression when disaggregated by sex. Data from the Allen Brain Institute.

**Supplementary Figure S5.**
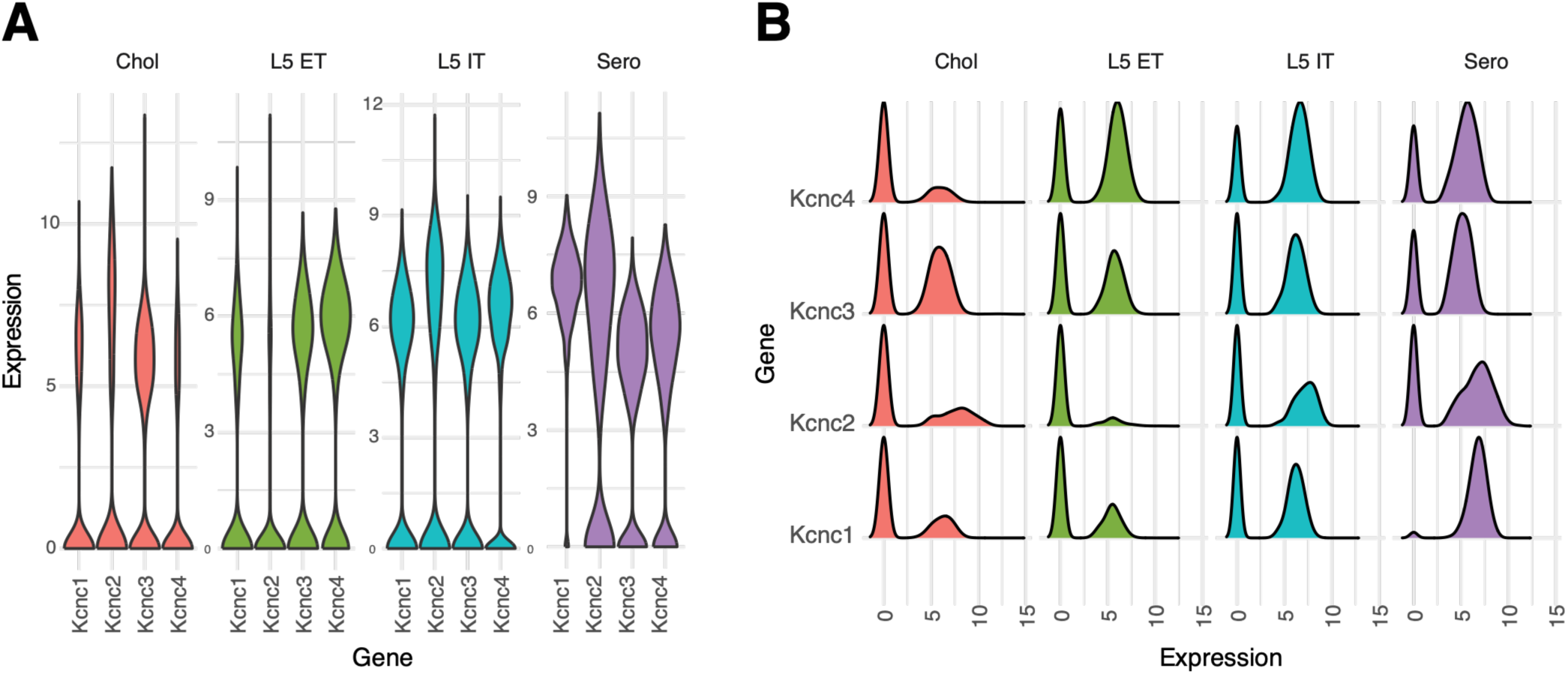
Expression of Shaw voltage-gated potassium channel subunits in some relevant cell classes. **A.** Violin plots and **B.** ridge plots of Shaw voltage-gated potassium channel subunit (Kv3) expression. Expression is visualized across cholinergic cells (red), two subtypes of cortical glutamatergic layer 5 cells (L5 ET, green; L5 IT, blue), and serotonergic cells (purple). Data from the Allen Brain Institute.

**Supplementary Figure S6.**
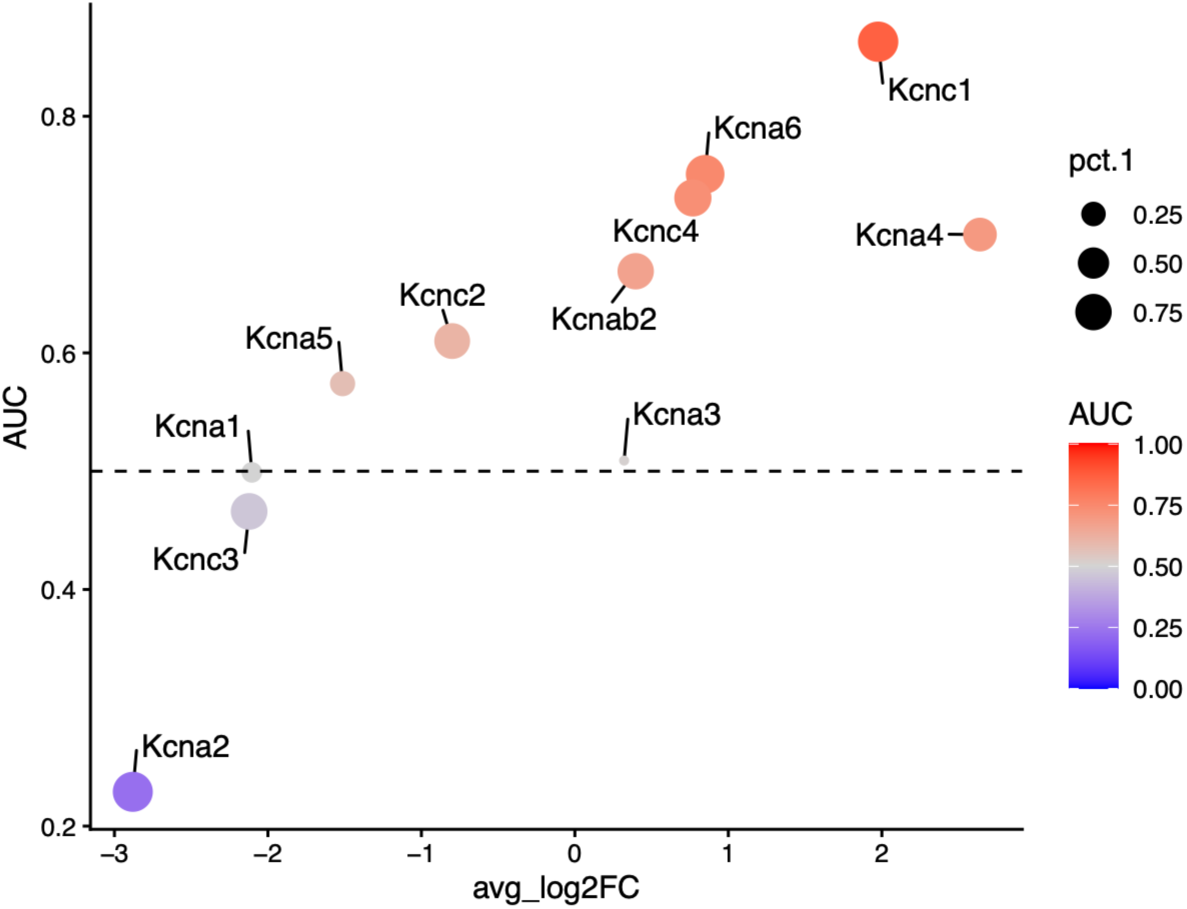
Differential expression of Shaker and Shaw voltage-gated potassium channel subunits between serotonergic and cholinergic cells. Dot plot depicting receiver operating characteristic (ROC) differential gene expression analysis of voltage-gated potassium channel (Shaker and Shaw) subunit expression between serotonergic and cholinergic neurons. Quantitative analysis probes the differences between serotonergic and cholinergic cells in their pattern of Shaker and Shaw subunit expression. Several subunits stand out in predicting neurotransmitter identity. The expression levels of Shaker subunits, Kv1.6 (*Kcna6*) and Kv1.4 (*Kcna4*), and Shaw subunits, Kv3.1 (*Kcnc1*) and Kv3.4 (*Kcnc4*) are more likely to predict serotonergic identity (AUC > 0.50). The expression levels of *Kcna2* and *Kcnc3* are more likely to predict cholinergic identity (AUC < 0.50). The diversity of subunit expression in serotonin neurons highlights the possibility of heteromultimeric channels in the Shaker and Shaw families. Data sourced from the Allen Brain Institute.

